# Role of Klhl14 in senescence and epithelial-to-mesenchymal transition *via* TGF-β modulation

**DOI:** 10.1101/2025.09.12.675818

**Authors:** Rufina Maturi, Abel Soto-Gamez, Anne L Jellema-de Bruin, Matteo Esposito, Schelto Kruijff, Gabriella De Vita, Rob P. Coppes

## Abstract

KLHL14, a component of an E3-ubiquitin ligase complex, has emerged as a context-dependent oncogene or tumor suppressor, particularly important for thyroid development. Yet its role in thyroid biology remains largely unexplored. In this study, we uncover a central function for KLHL14 in maintaining thyroid epithelial identity and regulating tissue homeostasis. Using a thyroid organoid model, we show that KLHL14 is essential for the proper growth and maturation of thyroid cells. Reduction of KLHL14 expression disrupts organoid development and triggers a dual cellular response involving features of both senescence and epithelial-to-mesenchymal transition. These phenotypic changes are accompanied by increased cellular plasticity and migratory capacity. Mechanistically, we identify TGF-β signaling as a key pathway activated upon KLHL14 depletion, contributing to the observed cellular reprogramming. Inhibiting TGF-β restores growth and reduces markers of senescence and EMT, positioning KLHL14 as an upstream modulator of this signaling axis. These findings reveal a previously unrecognized role for KLHL14, suggesting that its dysfunction may contribute to disease progression in aggressive thyroid cancers. This work broadens our understanding of thyroid epithelial biology and provides molecular insights extendable to other tissues, highlighting KLHL14 as a potential target for therapeutic interventions in malignancies displaying the herein explored features.

## Introduction

Kelch-like protein member 14 (KLHL14) is a member of the KLHL family, a major class of substrate adaptors for Cullin-RING E3 ligases involved in the ubiquitin-proteasome system (UPS)^1^. *Klhl14* and its antisense gene, the lncRNA *Klhl14-AS*, were found among the top 20 enriched transcripts in the E10.5 thyroid bud, together with the most important thyroid transcriptional factors, such as *Hhex* and *Pax8*^2^. Moreover, *Klhl14* silencing in healthy thyroid cells resulted in a drop in the expression of thyroid-specific markers, suggesting this protein plays a key role in maintaining cell differentiation^3^. Klhl14 expression is high specifically in the thyroid, B1a-lymphocytes, and spleen. Indeed, it is necessary to define B-cell differentiation and neoplastic transformation. Furthermore, a lack of *Klhl14*, even in heterozygosity, results in a severe deficiency of B1a-lymphocytes and an increase in B1b-lymphocytes^4^.

Recently, KLHL14 has been identified as a tumor suppressor in thyroid cancer, with its loss being proportional to the undifferentiated status of cancer cells. Additionally, the protein rescue was able to curb tumor cell proliferation and an apoptotic rate closer to those of healthy thyroid cells^3^.

KLHL14’s involvement in regulating proliferation and apoptosis might also be related to cellular senescence. Cellular senescence is a response to stress stimuli associated with a permanent cell cycle arrest and a heterogeneous plethora of morphological and functional changes^5^. Initially, senescence was identified as a mechanism to prevent the proliferation of damaged cells. This process is also implicated in aging, cancer, and various age-related diseases^6^. Senescence is considered an anti-tumor tool usually associated with a positive treatment response. Nevertheless, some cancer treatments, like radiotherapy or chemotherapy, may promote the accumulation of senescent tumor cells, which promote drug resistance and metastasis, especially via the senescence-associated secretory phenotype (SASP)^7^.

Beyond thyroid disorders, Klhl14 has been linked to several diseases, i.e., early-onset generalized dystonia, where the pathogenesis could be linked to the interruption of KLHL14 interaction with mutated TorsinA, a protein involved in the regulation of neuronal protein trafficking^8^. Moreover, Klhl14 has been described on one hand as a tumor promoter in ovarian and endometrial cancer, and on the other hand, as a tumor suppressor in diffuse large B cell lymphoma (DLBCL) and malignant mesothelioma^9–12^. Lack of KLHL14 promotes tumor proliferation, migration, and invasion, being reliant on TGF-β. Indeed, TGF-β turned out to promote KLHL14 *de novo* synthesis, stability, and activation of nuclear-cytoplasmic shuttling^12^.

The interplay between KLHL14 and TGF-β is particularly interesting considering TGF-β’s role in epithelial-to-mesenchymal transition (EMT) and the importance of this process in tumor progression^13^. EMT is a biological process in which polarized epithelial cells, usually bound and interacting with the basement membrane, through morphological and functional changes, transform into mesenchymal cells. The phenomenon is characterized by the loss of epithelial characteristics, such as cell-cell adhesion and polarity, and enhancement of mesenchymal traits, including increased migratory capacity, invasiveness, loss of polarity, and cell adhesion. EMT plays a crucial role in physiological and pathological events, like embryogenesis, tissue regeneration, cancer progression, and metastasis formation^14^.

Given Klhl14’s broad involvement in several tissues’ malignancies and developmental disorders, along with important findings in physiologic as well as neoplastic thyroid, in this study we aimed to investigate the molecular mechanisms through which it plays a role in thyroid development and carcinogenesis, exploiting tissue-derived organoids.

## Results

### 1. Klhl14 expression pattern overlaps with thyroid differentiation markers

Tissue-derived thyroid organoids provide a physiologically relevant model for investigating the role of specific proteins within a three-dimensional (3D) multicellular architecture that more accurately recapitulates the *in vivo* thyroid environment compared to conventional two-dimensional (2D) cultures^15^.

In this study, thyroid cells isolated from murine tissue were embedded in a basement membrane matrix (BMM) and cultured, as outlined in the experimental scheme, for 5, 7, and 11 days, (Fig. 1a). To first assess *Klhl14* gene expression in these cells, qPCR analysis on mouse thyroid gland organoids (mTGOs) collected at the three timepoints of culturing was performed. During organoid formation, single cells within the BMM proliferate and compose the 3D complex structure, potentially changing their transcriptomic profile. *Klhl14* mRNA levels (Fig. 1b) and protein levels (Fig. 1c) increased. Specifically, KLHL14 immunofluorescent staining was barely detectable on day 5, more pronounced on day 7, and even higher on day 11.

**Fig. 1:**
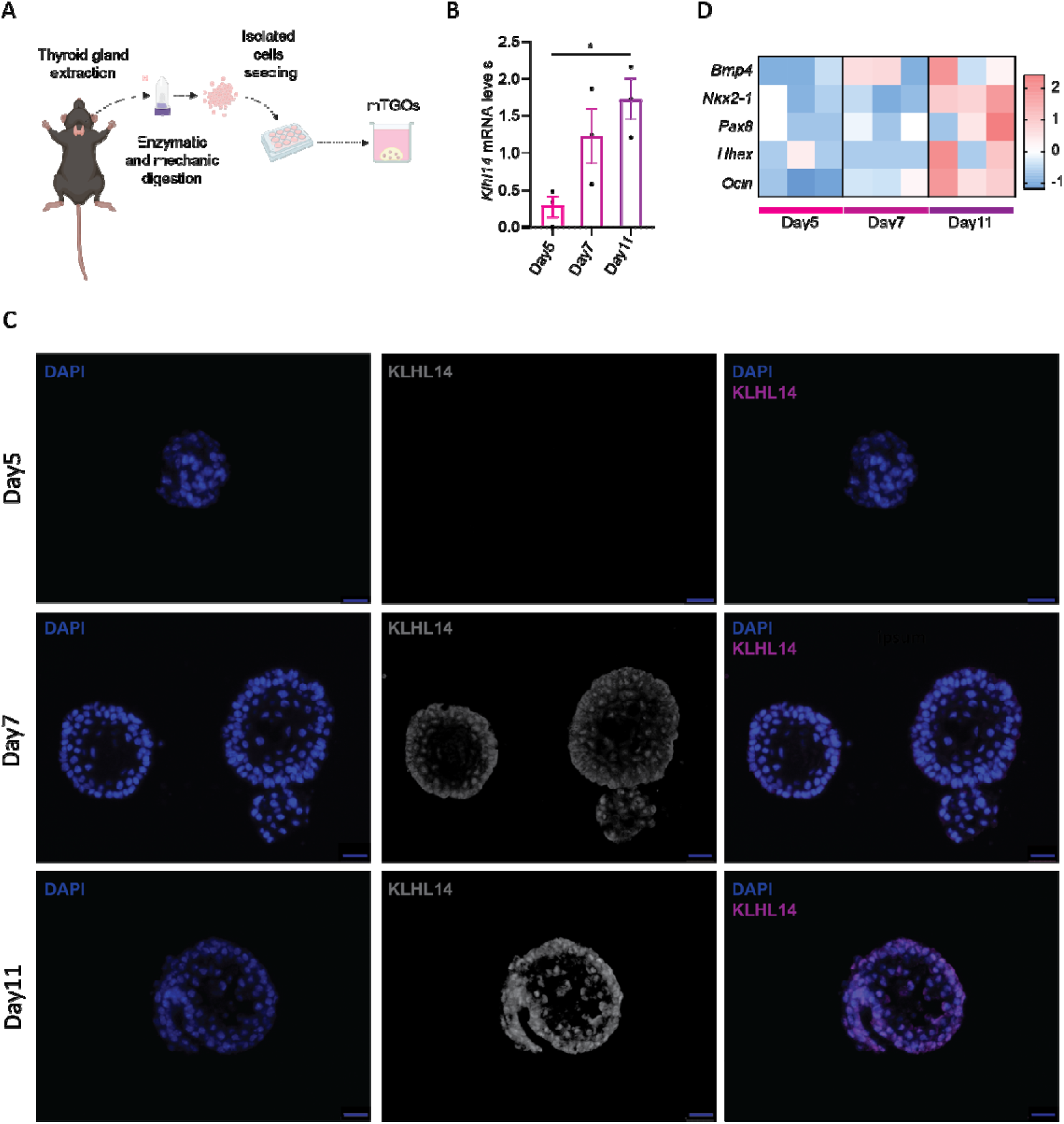
*Klhl14* expression pattern parallels thyroid differentiation markers in mTGOs. (**a**) General experimental setup for mTGO culture: cells are isolated from mice thyroid glands by enzymatic and mechanical digestion, then seeded in a solid BMM and cultured for 5, 7, or 11 days. (**b**) qPCR expression of *Klhl14* in 5-, 7-, and 11-days old organoids. (**c**) Immunofluorescent staining of KLHL14 protein levels in 5-, 7-, and 11-days old organoids. Scalebar 25 μm. (**d**) Bulk RNA sequencing of 5-, 7-, and 11-days old organoids. The heatmap represents the count-per-million Z-scores of the expression of genes related to thyroid maturation and differentiation at the three considered time points, highlighting how a higher definition of a thyroid profile accompanies the growth of the three-dimensional structure. Data are shown as mean ± SEM. Group differences were evaluated with two-tailed Student’s *t*-tests, n=3. \**p* < 0.05

Bulk RNA sequencing analysis revealed a progressive increase in the expression of key transcription factors involved in thyroid maturation, including *Bmp4, Nkx2-1, Pax8, Hhex*, and *Ocln* (Occludin), consistent with their established roles in thyroid cell lineage specification^16^ (Fig. 1d). Indeed, the heatmap visualization of Z-scores further illustrated a robust correlation between organoid growth and the induction of thyroid-specific gene expression.

Thus, it is interesting to note that *Klhl14* mRNA expression followed a trajectory parallel to that of classical thyroid differentiation markers, suggesting a potential role for *Klhl14* in thyroid maturation.

### 2. KLHL14 knockdown impairs thyroid organoid growth

To further investigate KLHL14’s role in thyroid development, KLHL14 gene expression in organoids was downregulated using a short hairpin RNA (shKlhl14) sequence and a Green Fluorescent Protein (GFP) tag to follow the lentivirus-transduced cells (Fig. 2a). On average, the extent of silencing was around 60%, as indicated by Western blot analyses (Fig. 2b). Representative images of 7-day-old organoids grown from GFP^+^ cells are shown in Fig. 2c. Notably, the shKlhl14-treated (KLHL14 KD) spheres are smaller than control shSCRA-treated ones, presumably due to pronounced growth impairment. An Incucyte Live-Cell Analysis system was employed to quantify the organoids’ size more accurately. The measurement of the green intensity is a preliminary analysis to validate the coverage by organoids in the well (Fig. 2d, left). The ratio between the green intensity and the number of objects can be considered an indirect measurement of the dimension of the organoids (Fig. 2d, right). Both parameters show a significant decrease in shKlhl14-treated organoids. After one week of culturing, organoids were counted, considering a minimum threshold of 50 mm to define maturity (Fig. 2e, left). A significant reduction in shKlhl14-treated secondary organoids forming efficiency (OFE) was observed. To assess overall cell proliferation, after enzymatic digestion of the organoids, the total number of cells was counted to establish the population doublings (Fig. 2e, right).

**Fig. 2:**
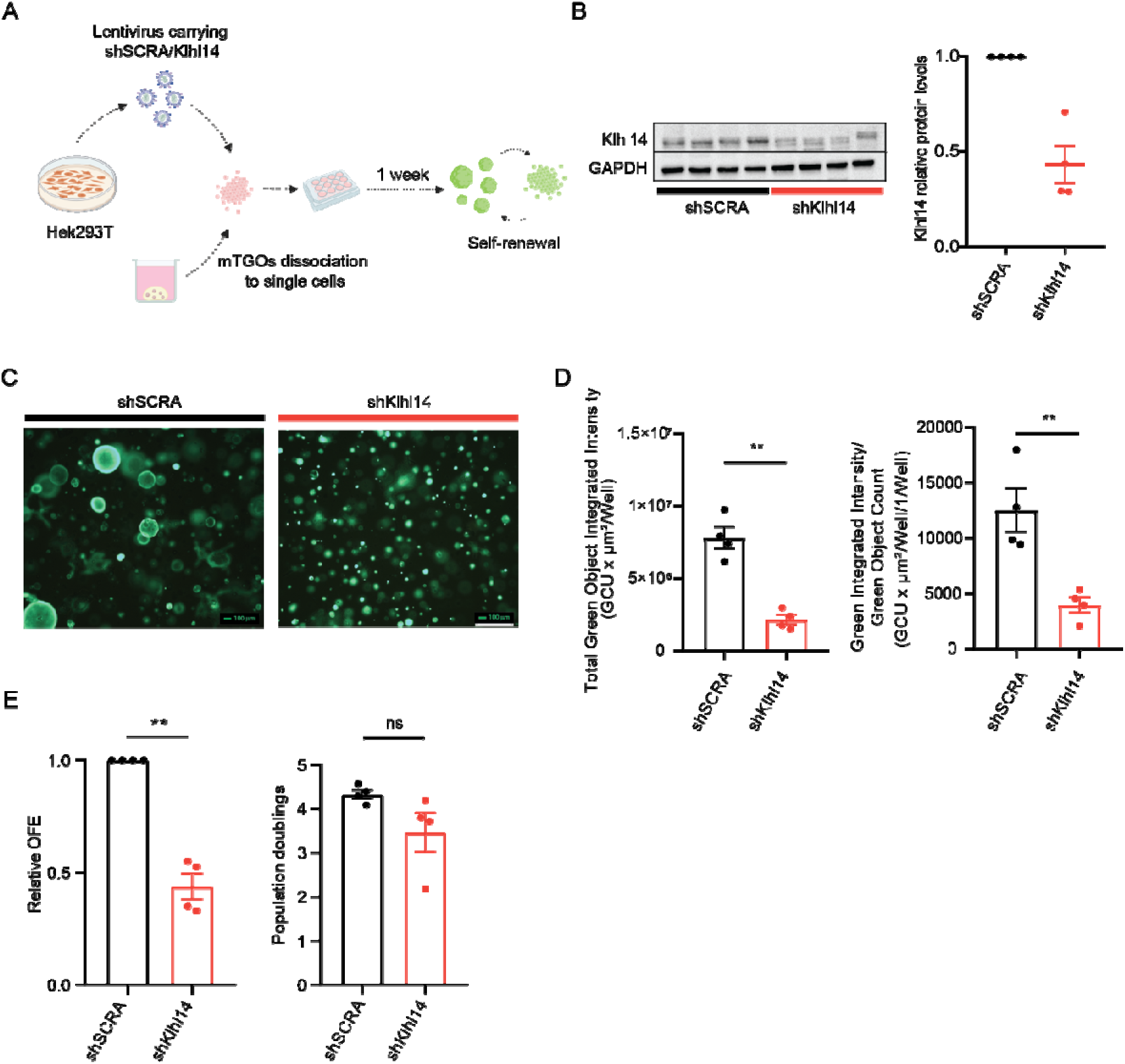
KLHL14 KD impairs thyroid organoid growth. (**a**) Experimental setup: single cells obtained from 7-day-old organoids were transduced with lentiviruses after 24 hours, seeded in BMM domes, and analyzed one week later. (**b**) Western blot analysis to validate KLHL14 silencing, using GAPDH as housekeeping. On the right is the relative normalized quantification of the bands. (**c**) Microscope images of organoids grown from transduced cells for one week, using the GFP channel, 4x objective. Scalebar 100 μm. (**d**) Data obtained from the Incucyte Live-Cell Analysis system exploiting the GFP expression of the organoids: on the left, the green integrated intensity. On the right, the ratio between the green integrated intensity and the total object count indicates the organoids’ dimension. (**e**) 7-day-old organoids obtained from transduced cells were collected after the disruption of the BMM domes, and relative OFE was calculated by counting organoids with a size > 50 mm (left graph). After trypsinization of the collected organoids, single cells were counted to calculate the population doublings (right graph) according to the formula described in the Materials and Methods. Data are shown as mean ± SEM. Group differences were evaluated with two-tailed Student’s *t*-tests. N=4. \*\**p* < 0.01

Notably, no significant decrease in cell number was detected, suggesting that Klhl14 silencing may be selectively detrimental to a small subset of cells. This discrepancy between the reduced OFE and the unchanged population doublings could be due to the threshold used for OFE analysis, meaning that many organoids failed to reach the minimum size cutoff of 50□μm.

These findings suggest the importance of *Klhl14* expression and its role in fine-tuning for thyroid cell proliferation, survival, maturation, and differentiation.

### 3. Proteomic analysis defines KLHL14 KD organoids’ molecular profile

A whole proteomic analysis was performed to explore the molecular features of KLHL14 KD organoids related to the development differences observed. Principal component analysis (PCA) revealed that Klhl14 KD samples cluster separately from the control samples, confirming a divergent proteomic profile associated with KLHL14 silencing. (Fig. 3a). As shown in Fig. 3b, 83 proteins were overexpressed, while 194 were significantly less abundant in the shKlhl14-treated group compared to the shSCRA-treated controls. To look at key players in the phenotype switch process, gene set enrichment analysis (GSEA) was performed, and the main dysregulated biological processes after KLHL14 KD were unveiled. Fig. 3c reports the top 10 gene ontology (GO) terms for up- and down-regulated proteins. Of note, among the most downregulated key processes are *epidermis development*, *negative regulation of response to external stimulus,* and *negative regulation of cytokine production*, suggesting impairment in several aspects: being underdeveloped, lacking the ability to face stress, and activating an unbridled inflammatory response. GO terms associated with the upregulated proteins are *apoptosis-related proteins* and *negative regulation of cell projection organization* involving key molecules such as Vimentin and *leukocyte proliferation* and *migration,* which comprise key factors such as CD44, P16, and P14ARF (Fig. 3b and d). Considering KLHL14 role as substrate-recognition subunit of ubiquitin ligase complexes, this study focused on the overexpressed proteins as a result of its knockdown. The related 15 most significant upregulated proteins in the shKlhl14 group are shown in Fig. 3d. The first is VIMENTIN (*Vim*), a type III intermediate filament protein widely recognized as a hallmark of mesenchymal cells and a key marker of epithelial-to-mesenchymal transition (EMT)^17^. GALECTIN-1 (*Lgals1*) is a lectin-binding beta-galactoside, was also markedly upregulated; it plays a multifaceted role in proliferation and apoptosis, capable of either promoting or inhibiting cell division depending on context^18^. Another prominent protein is CLUSTERIN (*Clu*), an extracellular molecular chaperone whose role is to prevent the aggregation of misfolded proteins. Its expression is notably induced by TGF-β signaling^19^, known to be a pivotal factor in the regulation of many biological processes, such as EMT^13^. Importin subunit a1 (*Kpna2*), a key component of the nucleocytoplasmic transport, was also significantly upregulated; its dysregulation is a major hit in many tumorigenic events^20^. P16 and P14ARF, two cyclin-dependent kinase inhibitors encoded by the same locus, *Cdkn2a*, were found to be elevated. These proteins oversee the progression through the cell cycle, and P16 is pivotal in the cell cycle arrest occurring with senescence^21^. Among the remaining significantly upregulated proteins, CD44 (*CD44*) is a cell adhesion molecule also functioning as a hyaluronic acid receptor, involved in a wide range of physiological and pathological processes, primarily used as a stem cell marker and EMT-related protein^22,23^. In parallel, apoptosis antagonizing transcription factor (*Aatf*) is a transcriptional coactivator correlated with cell cycle progression through all checkpoints, an apoptosis regulator, and is involved in the DNA damage response. Known for its eclectic role, it has been linked to many cancer types ^24^. Finally, Melanoma cell adhesion molecule (*Mcam*) is another interaction-mediating protein and a receptor for some growth factors; its expression can be regulated via TGF-β and increases during development but also metastatic cancer progression, being a main characteristic of EMT^25^. Collectively, all these data point towards cellular senescence and EMT as key changes that may underlie the growth impairment of shKlhl-treated organoids.

**Fig. 3:**
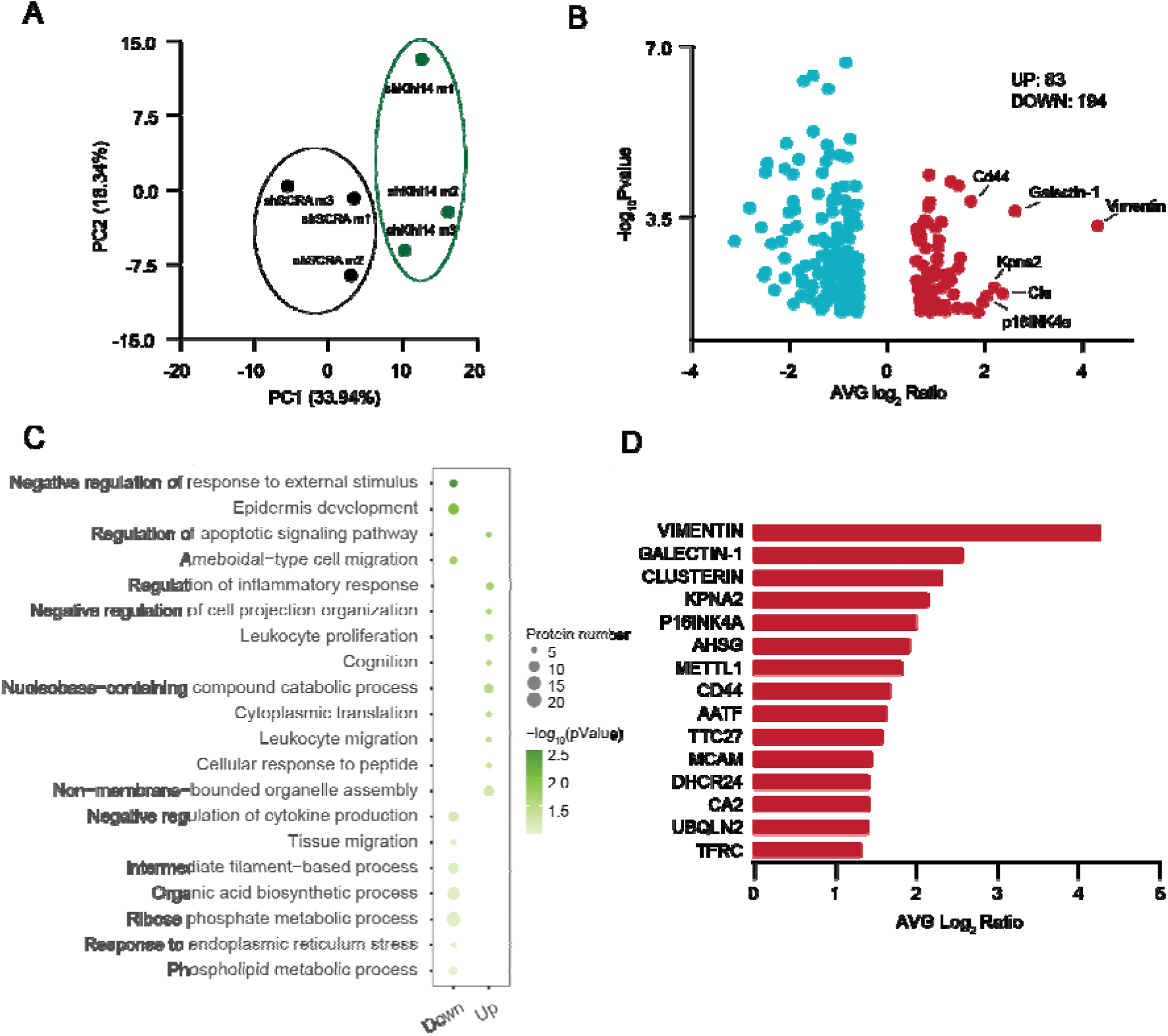
Proteomic analysis of KLHL14 KD organoids revealed key dysregulated pathways. (**a**) PCA plot of 7 days-old shSCRA and shKlhl14 treated organoids obtained from proteomic data. (**b**) Normalized results from the whole proteomic analysis were used to evaluate significantly dysregulated proteins in 7-day-old shKlhl14 organoids: 83 molecules are overrepresented, whereas 194 are downregulated. (**c**) GSEA indicated key processes associated with the two groups of differentially expressed proteins, depicted with their significance scores (green shades) and the number of proteins belonging to each term. (**d**) Top 15 overexpressed genes in Klhl14 KD organoids with their respective -AVG Log2 ratio. N=3

### 4. KLHL14 KD induces premature senescence and EMT

To validate the hypothesis that KLHL14 KD induces a phenotypic switch from epithelial to mesenchymal, the levels of the transcription factors *Snai1*, *Twist1,* and *Zeb1*, master regulators of the EMT, were assessed using qPCR. Interestingly, a significant increase was found for *Twist1* and *Zeb1* levels but not for *Snai1* (Fig. 4a). Moreover, an increase in the expression levels of mesenchymal markers such as *Vim* and *Cdh2* was found, further supporting the induction of EMT following KLHL14 silencing (Fig. 4b).

**Fig. 4:**
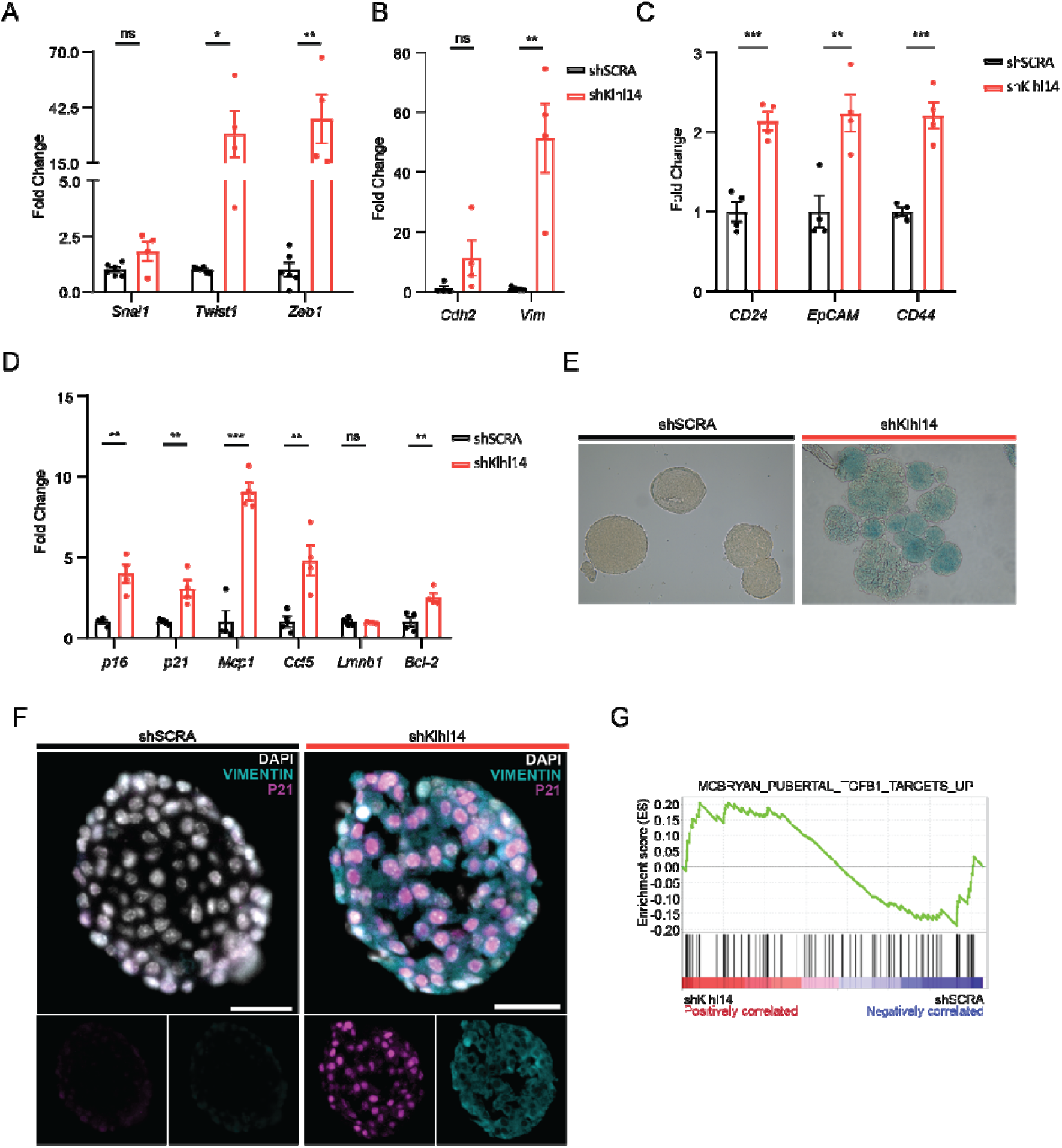
KLHL14 KD organoids show a senescence and EMT phenotype. (**a**) 7-day-old organoids obtained from transduced cells were collected and analyzed by qPCR to evaluate the expression levels of senescence markers in shKlhl14 organoids. Data are shown as fold change relative to shSCRA organoids. (**b**) Representative images of shSCRA and shKlhl14 organoids after beta-galactosidase staining performed on 7-day-old organoids: the reaction’s product gives the blue signal reflecting the enzyme’s activity. (**c**) qPCR analysis on 7-day-old organoids obtained from transduced cells evaluating the expression levels of EMT-related genes: *Snai1*, *Twist1,* and *Zeb1* as key transcriptional factors. (**d**) qPCR analysis on 7-day-old organoids obtained from transduced cells evaluating the expression levels of *Cdh2* and *Vim* as EMT effectors molecules. (**e**) Gene expression analysis through qPCR for *CD24*, *EpCAM,* and *CD44* was performed on shSCRA and shKlhl14 7-day-old organoids to explore the dedifferentiation process activated with the knockdown. (**f**) GSEA of the proteomic analysis data unveiled the TGF-β signaling pathway activation in shKlhl14 organoids compared to the controls. Data are represented considering the ES given by the software. (**g**) 7-day-old organoids grown from transduced cells were fixed and embedded to perform immunofluorescence staining for P21 and VIMENTIN. Scalebar 25 μm. Data are shown as mean ± SEM. Group differences were evaluated with two-tailed Student’s *t*-tests. N=4. \**p* < 0.05, \*\**p* < 0.01, \*\*\**p* < 0.001

Next, the occurrence of senescence was confirmed for many senescence-associated genes by qPCR analysis. The upregulated expression of *p16* and *p21* transcripts indicated an extensive cell cycle arrest in shKlhl14 samples, paralleled by enhanced production of SASPs, including *Ccl5* and *Mcp1*, as well as elevated levels of the anti-apoptotic gene Bcl-2. In contrast, the expression of *Lmnb1*, a gene typically downregulated during senescence, remained unchanged (Fig. 4d). To further validate this, a SAS-β-Gal staining was performed. Figure 4e shows representative images of the two conditions: shSCRA with big, clean organoids, where a blue signal from β-Gal activity is barely visible. In contrast, shKlhl14 organoids are smaller, and the blue staining is intense, especially in the structures’ core, indicating cellular senescence.

Given the importance of KLHL14 and Klhl14-AS in the maintenance of thyroid cell differentiation^3,26^ and the results of proteomic analysis, we measured the expression levels of some stemness markers: *CD24*, *EpCAM,* and *CD44*^27,28^. These three glycoproteins have been described as (cancer) stem cell markers, playing a main role in cell adhesion and migration. Interestingly, all three were upregulated in KLHL14 KD organoids, supporting a KLHL14 silencing-associated de-differentiated phenotype (Fig. 4c).

Finally, to check whether senescent cells are the same ones undergoing EMT, co-staining of shSCRA/shKlhl14 organoids for VIMENTIN and P21 was performed. The representative image in Fig. 4f (right) shows a diffuse staining of both markers overlapping in most cells, suggesting the presence of a relatively homogeneous population of senescent cells going through EMT rather than two different populations. In contrast, shSCRA-treated organoids exhibited only weak staining for both markers, proving that, in normal conditions, both markers are expressed only at very low levels.

Transforming Growth Factor (TGF) is known to be involved in all the processes described above, as suggested for malignant mesothelioma^12^, and may also play a role in the KLHL14 KD phenotype. Indeed, GSEA analysis of the proteomics data from shKlhl14 organoids showed a positive enrichment score (ES) for the shKlhl14 organoids with the set of TGF-β target genes. On the other hand, a negative ES for the shSCRA samples was observed, highlighting an increase in the overall abundance of proteins activated downstream of the TGF-β signaling pathway when KLHL14 was knocked down (Fig. 4g).

To investigate the role of TGF-β in the development of mTGOs, they were treated with increasing concentrations of TGF-β and cultured for one week. This resulted in a severe impairment of organoid growth and population doublings, already at the lowest concentration (Supplementary Fig. 1a-c). Moreover, western blot analysis showed that even at a low concentration (0.01 ng/ml), and more noticeably at a higher concentration (0.05 ng/ml), TGF-β affects KLHL14. This effect does not appear to alter the total protein level (Supplementary Fig. 1d) but instead suggests posttranslational modifications (Supplementary Fig. 1e). Specifically, at 0.05 ng/ml, the KLHL14 band seen in the controls splits into two fainter bands of similar molecular weight, suggesting that TGF-β may influence KLHL14 homeostasis or stability through posttranslational modification. These data show that KLHL14 KD enhances TGF-β signaling, which may activate the EMT and senescence programs, leading to severe growth impairment. In turn, TGF-β has an effect on KLHL14 at the protein level.

### 5. TGF-β-inhibition rescues KLHL14 KD phenotype

To further substantiate the role of TGF-β in KLHL14 KD mTGO, we pharmacologically inhibited TGF-β receptors using A83-01 (Tocris), a selective inhibitor targeting the type I TGF-β receptor ALK5 kinase, as well as the type I nodal receptors ALK4 and ALK7. The experimental setup is illustrated in Fig. 5a. After the shSCRA or shKlhl14 lentiviral transduction, cells were seeded with the drug (or DMSO as control) and analyzed seven days later, using qPCR and immunofluorescence. Representative images of the four conditions show green fluorescence, marking only organoids derived from transduced cells. The images demonstrate that TGF-β inhibition restores shKlhl14 organoids morphology, forming larger structures (Fig. 5b). Consistently, both OFE and population doublings were normalized in the shKlhl14+/TGF-β inhibitor (TGF-βi) group, reaching levels comparable to shSCRA controls, indicating an increased fitness of shKlhl14+TGF-βi (Fig. 5c and 5d). Moreover, qPCR analysis showed reduced EMT-driving transcriptional factors, confirming that the TGF-βi counteracts the increase in *Twist1* and *Zeb1* levels induced by shKlhl14 (Fig. 5e). Similarly, mesenchymal markers such as *Cdh2* and *Vim* were comparable to shSCRA samples (Fig. 5f).

**Fig. 5:**
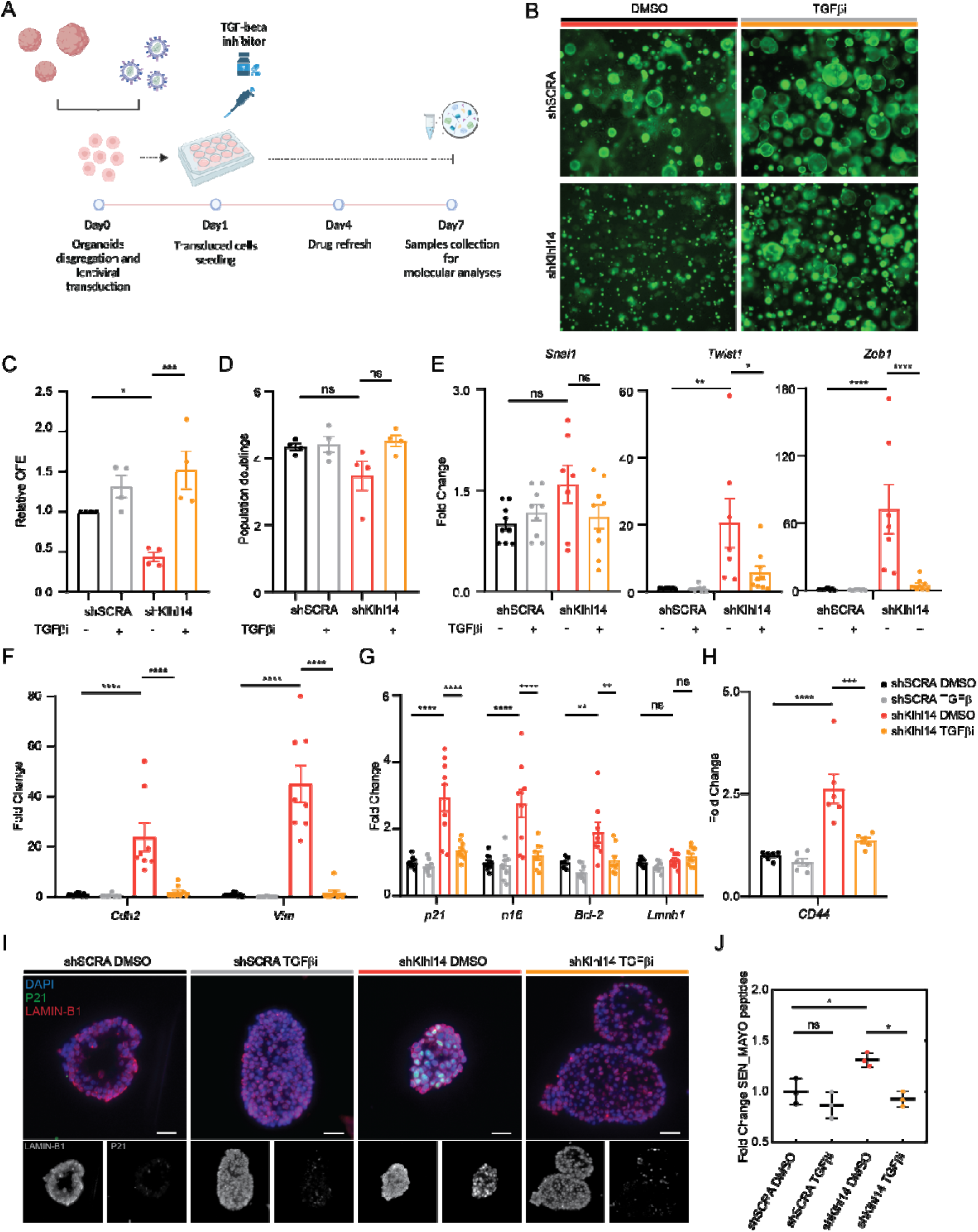
TGF-β-inhibition rescues KLHL14 KD-induced phenotype. (**a**) Experimental set-up scheme: mature organoids were used to obtain single cells to transduce with a lentiviral vector carrying a shKlhl14 RNA sequence (or a shSCRA for controls). 24 hours later, cells were counted and seeded with TGF-βi (or DMSO as control) and analyzed 7 days later. (**b**) Representative microscope images of organoids grown from transduced cells with TGF-βi or DMSO after one week, using the green fluorescence channel, 4x objective. (**c**) 7-day-old organoids grown with TGF-βi (or DMSO) were collected after the disruption of BMM domes and counted (> 50 mm). Relative OFE was calculated for each sample related to its shSCRA DMSO control. N=4 (**d**) Dispersed organoid cells were counted to calculate the population doublings on day 7. N=4 (**e**) 7-day-old organoids grown with TGF-βi (or DMSO as control) were collected and analyzed using qPCR to assess the expression levels of EMT transcriptional factors. Data are presented as fold change relative to shSCRA DMSO samples. (**f**) qPCR analysis of EMT effector molecules presented as fold change relative to shSCRA DMSO samples. (**g**) qPCR analysis of senescence markers presented as fold change relative to shSCRA DMSO samples. (**h**) qPCR analysis of CD44 plotted as fold change relative to shSCRA DMSO samples. (**i**) 7-day-old organoids grown with TGF-βi (or DMSO as control) were fixed and embedded to perform immunofluorescence staining for P21 and LAMIN-B1. Scalebar 25 μm. (**j**) Average fold change of protein levels of SEN_MAYO gene set in shSCRA and shKlhl14 organoids with and without TGF-βi treatment. Data are shown as mean ± SEM. Group differences were evaluated with one-way ANOVA with post-hoc Tukey’s HSD. N=4-9 \**p* < 0.05, \*\**p* < 0.01, \*\*\**p* < 0.001, \*\*\*\**p* <0.0001

qPCR also indicated that shKlhl14 mTGOs grown in the presence of TGF-βi show normal/control levels of *p21*, *p16*, and *Bcl-2*. These data suggest that TGF-βi helps the cells avoid the expression of cell cycle arrest-associated factors and retain the organoids’ growth potential (Fig. 5g). Next, the expression of (cancer) stemness markers following TGF-βi was evaluated. Interestingly, *CD44* expression falls back to the control levels, meaning that the dedifferentiation process may also be counteracted by blocking TGF-β signaling (Fig. 5h). Moreover, co-staining for P21 and LAMIN-B1 confirmed a reduction of cellular senescence (Fig. 5i). Control organoids displayed no significant changes in the staining patterns, whereas the shKlhl14 DMSO showed many P21-positive (P21^+^) nuclei, with a lower LAMIN-B1 (LAMIN-B1^low^) signal. When treating shKlhl14 with TGF-βi, the number of P21^+^/LAMIN-B1^low^ nuclei decreases to control levels (Fig. 5i).

Next, proteomic analysis was performed on TGF-βi treated organoids, and results were exploited to analyze protein levels of the gene set SEN_MAYO, which includes a comprehensive panel of senescence-related factors (Fig. 5j). ShKlhl14 DMSO samples show a higher expression of this whole gene set than the shSCRA samples, whereas shKlhl14 TGF-βi were back to controls, indicating that the senescence program is reverted at the molecular level.

To investigate whether this mechanism is conserved in human cells, human thyroid gland organoids (hTGOs) were generated from patient-derived biopsies. The experimental procedure (Fig. 6a) closely mirrored that of the murine model, with the main difference being a longer culture period (three weeks) required for hTGO growth. An *ad hoc* lentiviral vector was packaged to silence Klhl14 in hTGOs, and transduced cells were grown in the presence or absence of TGF-βi (Fig. 6b). hTGOs shKlhl14 without any TGF-βi treatment were co-stained for P21 and Vimentin to assess the occurrence, respectively, of senescence and EMT to validate the results obtained in mTGOs. As observed in the murine model, the shKlhl14 hTGOs showed many P21^+^ nuclei and displayed a high expression of Vimentin compared to controls, confirming the induction of both senescence and EMT (Fig. 6c). The four conditions parallel the murine observed phenotypes, resulting in smaller shKlhl14 organoids, which can grow bigger with the addition of TGF-βi. The Incucyte Live-Cell imaging confirmed the phenotype studied and extended these findings to the human system, with a reduction in the organoid size and number, supporting the idea that Klhl14 may play a conserved role in regulating thyroid cell behavior across species (Fig. 6d and 6e).

**Fig. 6:**
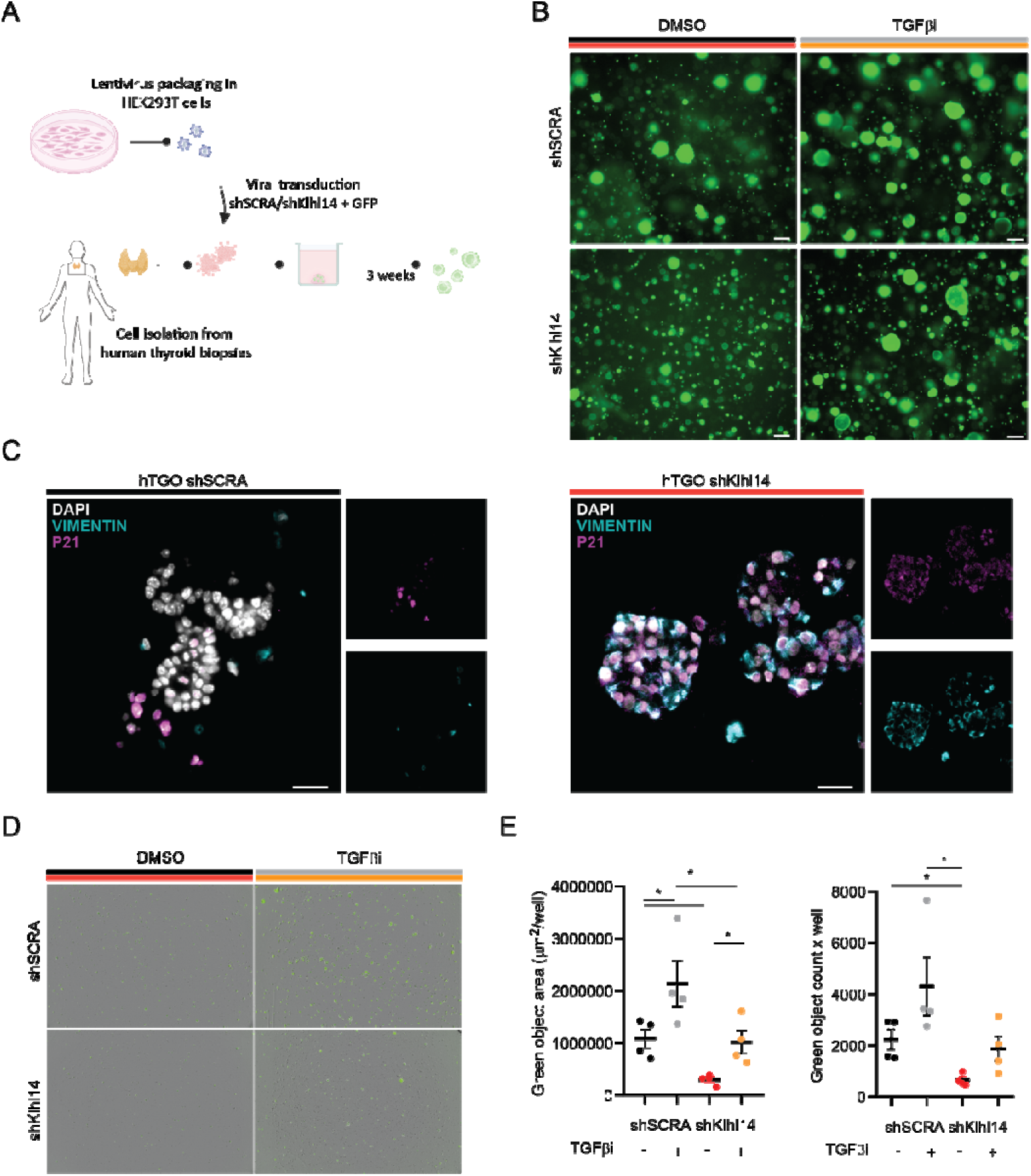
shKlhl14 hTGOs display the same phenotype as shKlhl14 mTGOs. (**a**) Experimental setup: single cells obtained from human thyroid biopsies were employed to grow hTGOs. Single cells derived from hTGOs disruption were transduced with lentiviruses after 24 hours, seeded in BMM domes, and analyzed three weeks later. (**b**) Microscope images of organoids grown from transduced cells for three weeks, using the GFP channel, 4x objective. (**c**) 21-day-old organoids grown from transduced cells were fixed and embedded to perform immunofluorescence staining for P21 and VIMENTIN. Scalebar 25 μm (**d**) Representative images acquired from the Incucyte Live-Cell Analysis system. (**e**) Data obtained from the Incucyte Live-Cell Analysis system exploiting the GFP expression of the organoids: the green object area and green object count. Data are shown as mean ± SEM. Group differences were evaluated with one-way ANOVA with post-hoc Tukey’s HSD. N=4. \**p* < 0.05

Collectively, these results suggest that Klhl14 KD triggers both EMT and senescence *via* TGFβ enhancement, and that these mechanisms are conserved between mouse and human thyroid organoids.

### 6. KLHL14 KD induces a switch in adhesion molecules

To further investigate the phenotype of mTGO, all the adhesion molecules detected during the whole proteomic analysis were classified either in epithelial or mesenchymal, and their levels (plotted as Z scores) were compared (Fig. 7a). The heatmap shows that epithelial adhesion molecules are downregulated in shKlhl14 DMSO samples to rescue levels comparable to the controls when adding the TGF-βi. In contrast, mesenchymal adhesion markers showed the opposite trend (Fig. 7a). To assess whether the mesenchymal phenotype was also responsible for the higher motility of shKlhl14 cells, organoids were disrupted, and single cells from shSCRA and shKlhl14 organoids were seeded on a BMM-coated plate to perform a scratch assay, with or without TGF-βi. Images obtained with the fluorescence microscope revealed that shKlhl14 could not attach to the coated plate. Therefore, it was not possible to make the scratch; instead, with the addition of TGF-βi in the media, shKlhl14 cells adhered (Fig. 7b). Trypan blue staining confirmed that shKlhl14 floating cells were still alive (Fig. 7c). Finally, 7-day-old organoids were seeded in 2D on a BMM-coated plate, and 24 hours later, the Incucyte Live-Cell imaging system was used to evaluate whether shKlhl14 organoids presented more migrating cells than the controls. In Fig. 7d (left), the squares represent organoids with (red square) or without (yellow square) migrating cells; the graph shows the percentage of organoids with migrating cells for each condition (Fig. 7d, right). This last data finally confirms that shKlhl14 organoids present more migrating cells than the controls, which is consistent with the induction of an EMT phenotype.

**Fig. 7:**
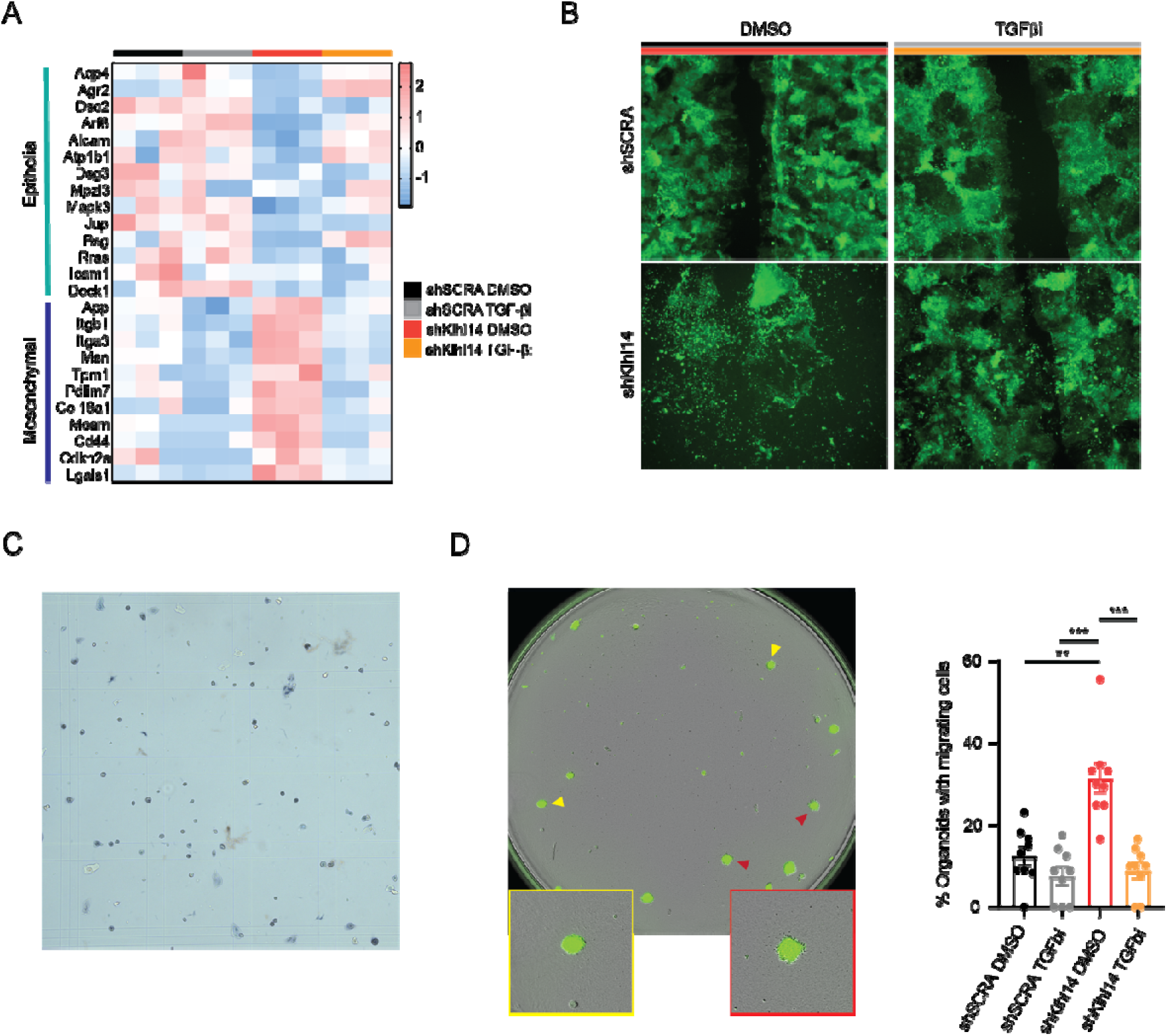
KLHL14 KD induces an adhesion molecule switch and enhanced migration. (**a**) Z-scores of adhesion molecules retrieved from whole proteomic analysis in 7-day-old shSCRA and shKlhl14 organoids, with and without TGF-βi treatment. (**b**) Representative microscope images of transduced cells seeded in 2D on a BMM-coated plate, with TGF-βi or DMSO after 24 hours, using the green fluorescence channel, 4x objective. (**c**) Light microscope image of trypan blue assay on shKlhl14 DMSO cells grown in 2D. (**d**) (left) Sample image acquired by the Incucyte Live-Cell imaging system: in the yellow box is an example of organoids without migrating cells; in the red box is an example of an organoid with migrating cells. (right) Percentage of organoids with migrating cells per each condition. N=3 n=3 ± SEM. Group differences were evaluated with one-way ANOVA with post-hoc Tukey’s HSD. \*\**p* < 0.01, \*\*\**p* < 0.001

## Discussion

This study unveils for the first time a role of KLHL14 as a protein that regulates senescence and EMT through fine-tuning of the TGF-β pathway activation in the thyroid. To date, KLHL14, a subunit of an E3-ubiquitin ligase complex, has been implicated in several biological processes, ranging from proteostasis to tumorigenesis. Its early activation during thyroid embryonic development (E10.5) suggests a central role for this protein in thyroid homeostasis. Moreover, it has emerged as a context-dependent oncogene or tumor suppressor, depending on the tissue^3,9,11,12^. For instance, in anaplastic thyroid carcinoma, it has been demonstrated that downregulation of KLHL14 promotes proliferation and survival^3^. Still, the molecular mechanisms underlying its function remain largely undefined.

This study utilized a tissue-derived thyroid organoid model^15^ to investigate the role of KLHL14 in thyroid physiology. Notably, Klhl14 expression increased during organoid growth, reflecting the dynamics of key thyroid differentiation regulators such as Hhex, Nkx2-1, Pax8, and Bmp4^29^, suggesting a role for KLHL14 in thyroid maturation. Similar findings were obtained for Klhl14 antisense lncRNA (Klhl14-AS), which modulates thyroid-specific gene expression via Bcl-2/Pax8 ceRNA circuits^26^. The sense-antisense gene pair indeed often exhibits cross-regulated behavior, suggesting the existence of a complex regulatory mechanism between the two.

Silencing of KLHL14 led to impaired thyroid organoid growth and viability, shedding light on the involvement of this protein in two biological processes that are pivotal in development and cancer biology. The importance of KLHL14 has been previously underlined when neonatal lethality was observed in Klhl14-*null* mice^4^. Proteomic profiling of KLHL14 KD organoids revealed extensive molecular rewiring and directed this study towards two biological processes, which may have been responsible for the observed phenotypes: EMT and cellular senescence. Following molecular analyses, it was confirmed that KLHL14 depletion induced signatures of both. Precisely, master regulators of cell cycle arrest (p16, p21) and SASPs (e.g., Ccl5, Mcp1) were upregulated, alongside mesenchymal markers (Vim, Cdh2) and EMT transcription factors (Zeb1, Twist1). EMT and senescence occurred concurrently within the same cells, rather than in different subpopulations: as such, an unusual form of *aberrant EMT* associated with cell cycle arrest and inflammatory phenotype was activated. While this is the first evidence of KLHL14 involvement in senescence, indicating a new biological function of the protein related to cell cycle regulation, its relevance in EMT is not new. Indeed, an *in vitro* study using amniotic epithelial cells as a spontaneous model of EMT revealed that it is the most downregulated gene in mesenchymal cells compared to epithelial ones^30^. Additionally, in malignant mesothelioma, Klhl14 expression has been shown to prevent EMT, as its downregulation is associated with the activation of this biological process, thereby contributing to cancer progression. In pituitary neuroendocrine tumors, the translocation of KLHL14 from the cytoplasm to the nucleus, together with E-cadherin mislocalization, has been associated with increased invasiveness, again suggesting the activation of EMT^31^.

Furthermore, herein, KLHL14 KD induced the expression of (cancer) stemness markers (CD24, CD44, EpCAM), indicating KLHL14 as a candidate for the maintenance of thyroid cell differentiation, but also supporting the occurrence of EMT^27,28^. Indeed, this study is consistent with previous findings associating late-stage thyroid and bladder cancers with the downregulation of KLHL14 and increased metastatic potential^3,32^.

Interestingly, proteomic analyses and functional experiments implicated TGF-β signalling as a key mediator of the phenotype observed upon KLHL14 depletion. The implication of TGF-β in thyroid follicular cell migration and follicle neogenesis was outlined decades ago, when porcine thyroid follicles or primary cultures were exposed to TGF-β in combination with EGF^33,34^. Furthermore, a recent study demonstrated that TGF-β induces EMT through SNAI1 modulation in thyroid cancer cells^35^. Therefore, our findings on the interplay between KLHL14 and TGF-β depict a new central molecule for thyroid physiology and potentially cancer development.

KLHL14 KD in organoids activates TGF-β downstream pathways, causing cells to undergo a complete switch in their adhesion molecules’ expression, displaying a new set of mesenchymal cell-cell and cell-matrix interaction proteins. Pharmacological inhibition of TGF-β signalling rescues growth impairment, reducing senescence and EMT markers.

Furthermore, a recent study in salivary gland organoids associated high CD44 levels with accentuated migratory capacity of progenitor cells^36^. This finding corroborates the results reported in this study: KLHL14 KD increases CD44 levels and leads to an enhanced migratory ability of thyroid cells, following the activation of TGF-β signaling.

Finally, preliminary experiments performed with hTGOs revealed that the KLHL14 KD shares the phenotype with mTGOs, paving the way to unveil a crucial and conserved mechanism of the KLHL14-TGF-β interplay.

These results establish, for the first time, KLHL14 as an upstream modulator of TGF-β activity, building on previous findings that showed TGF-β regulates KLHL14 stability and localization in malignant mesothelioma^12^.

Finally, our data reveal a regulatory role for TGF-β in inducing senescence, consistent with reports that describe its modulation of cyclin-dependent kinase inhibitors, ROS production, and telomerase activity^37–40^.

Despite the importance of the molecular findings of this work, the lack of *in vivo* validation restricts this knowledge to an *ex vivo* system in which immune components and systemic influences are absent. In a complete organism, there might be adaptation mechanisms activated to limit the events occurring after the downregulation of KLHL14. Moreover, to translate the results of this study to a clinical application exploiting Klhl14 as a target, further investigation to verify the occurrence of these mechanisms in cancerous models, like thyroid cancer organoids or mouse models, is needed.

In summary, we identified a crucial role for KLHL14 in thyroid organoid development through regulating proliferation, differentiation, EMT, and senescence. Our results highlight TGF-β as a key downstream effector of KLHL14 function, suggesting a bidirectional regulatory interaction between these molecules. This is the first time a specific molecular mechanism has been described, explaining the lethal phenotype observed in Klhl14-*null* mice^4^ and the link with aggressiveness observed in several tumor types. These insights advance our understanding of thyroid biology and suggest broader implications for KLHL14’s tumor suppressor functions across epithelial tissues.

## Materials and methods

### Isolation of murine thyroid gland cells and organoids culture

The animal procedures conducted in this study were approved by the Ethical Committee for Animal Testing at the University of Groningen. Thyroid glands from 8-12-week-old female C57BL/6 mice (Harlan, The Netherlands) were utilized. Thyroid tissues from three mice were pooled in a single tube containing Hank’s Balanced Salt Solution (HBSS) with 1% BSA. The tissue was mechanically dissociated using the GentleMACS dissociator (Miltenyi Biotech), followed by enzymatic digestion in HBSS/1% BSA buffer with CaCl_2_ (6.25 mM), collagenase I (100 U/ml; Gibco) and dispase (1.5 U/ml; Gibco). Cells obtained were expanded overnight in suspension in DMEM-F12 (Gibco), 1% penicillin/streptomycin (Gibco), GlutaMAX (2 mM; Gibco), epidermal growth factor (EGF) (20 ng/ml; Sigma Aldrich), fibroblast growth factor-2 (FGF-2) (20 ng/ml; Peprotech), and B27 supplement (0.5%; Gibco). After 1 day, cell suspension was collected, spheres dissociated into single cells using trypsin EDTA (0.05%). After filtering, cells were resuspended in DMEM-F12 (Gibco), counted and seeded: 25 µL of media containing 10.000 cells were combined on ice with 50 µL of Basement Membrane Matrigel (BMM; BD Biosciences) and deposited in the center of 12-well tissue culture plates. After solidification of the gel in the incubator for 30 min, 1 ml of thyroid growth medium (TGM), which comprised DMEM-F12 (Gibco), 1% penicillin/streptomycin (Gibco), GlutaMAX (2 mM; Gibco), epidermal growth factor (20 ng/ml; Sigma Aldrich), ROCK inhibitor Y-27632 (10 µM; Abcam, Cambridge, UK), fibroblast growth factor-2 (20 ng/ml; Peprotech), and B27 supplement (0.5%; Gibco).

### Collection and isolation of human thyroid glands

Non-malignant human thyroid tissue was obtained from donors aged 19–80 years undergoing scheduled thyroidectomy, following informed consent and approval by the institutional review board (Thyrostem study; METc 215/101). Tissue was collected from macroscopically normal regions of the thyroid, either contralateral to the tumor in patients with differentiated thyroid carcinoma (DTC) or from patients with goitre, a condition generally regarded as benign with a low risk of malignancy. All surgical resections were performed in accordance with clinical treatment plans approved by the institutional multidisciplinary thyroid board. Thyroid specimens were collected under sterile conditions and transported in HBSS (Gibco) supplemented with 1% BSA in Greiner tubes. Tissue was subjected to mechanical dissociation using the GentleMACS Dissociator (Miltenyi Biotec), followed by enzymatic digestion in HBSS/1% BSA containing collagenase I (100 U/mL; Gibco) and dispase (1.5 U/mL; Gibco). Following digestion, cells were seeded at a density of 4 × 10□ cells/well into a 24-well plate embedded in Basement Membrane Matrigel (BD Biosciences, 354234). Cultures were maintained in thyroid growth medium supplemented with Wnt and R-spondin (TGM+WR), consisting of DMEM/F12 containing, 10% UltraGROTM-PURE GI (Pelobiotech), 1% penicillin/streptomycin (Gibco), 2 mM GlutaMAX (Gibco), R-Spondin 1 (10 ng/ml; Sigma-Aldrich), EGF (20 ng/mL; Sigma-Aldrich), FGF-2 (20 ng/mL; Peprotech), 0.5% B27 supplement (Gibco), 1% heparin sodium salt solution (StemCell Technologies), nicotinamide (10 mM; Sigma-Aldrich), A83-01 (500 nM; Tocris Bioscience), and noggin (25 ng/mL; Peprotech).

### Self-renewal of thyroid organoids

One week (mouse) or three weeks (human) after seeding, the secondary spheres were counted, and the percentage of sphere-forming cells was calculated. We passaged these secondary spheres every time by first adding dispase (1 mg/ml) to the gels to release the spheres from them, then disrupted the spheres into single cells using trypsin-EDTA (0.05%) and then reseeding 10.000 single cells (mouse) or 25.000 (human) in Basement Membrane Matrigel, as described in the previous section. Organoid and single cell numbers were noted and used to calculate the organoid formation efficiency (OFE) and population doublings (PD) as indicated:

OFE%= (total number of organoids harvested / number of seeded cells) x 100

PD = ln (total number of harvested cells/number of seeded cells)/ln2

### Lentivirus packaging and transduction

For lentiviral production, HEK293T cells were cultured in DMEM supplemented with 10% FBS, Pen/Strep, and GlutaMAX. The cells were transfected with 3 µg of the vector of interest (pLV[shRNA]-EGFP:T2A>[shRNA]) (Vector Builder), 3 µg of the packaging plasmid (PAX2), 0.7 µg of the glycoprotein envelope plasmid (VSV-G), and 40 µL of polyethylenimine (PEI; 1 µg/mL). Two different lentiviruses were packaged: one to silence Klhl14 in mTGOs and one for hTGOs. The following day, the medium was replaced with DMEM-F12, and the viral supernatant was collected after 24 hours. For transduction, thyroid gland organoids between passage 2 and 5 were removed from the gels and dissociated into single cells. After counting, the cell suspension was incubated with viral supernatant and polybrene (6 µg/mL) in a 24-well plate overnight at 37°C with 5% CO2. The next day, the cells were counted and embedded in Basement Membrane Matrigel in a 12-well plate containing TGM. Three days later, the medium was supplemented with puromycin (1.5 µg/mL) for the selection of transduced cells, with media replacement every two days. Transduction efficiency exceeded 95% across all conditions and was evaluated by measuring the percentage of GFP-positive cells through the Quanteon flow cytometer at the Flow Cytometry Unit of the UMCG. Data was analyzed using the FlowJo software (v10.7.2).

### TGF-β treatment

Mouse thyroid organoid cells obtained from spheres trypsinization (trypsin-EDTA 0.05%) were seeded in Basement Membrane Matrigel as described in the *Self-renewal of thyroid organoids* section. Recombinant mouse TGF-β1 (Biolegend, 763102) was added to the culturing media to have 0.01 and 0.05 ng/ml as final concentration.

### TGF-βi treatment

To assess the role of TGF-β signaling on thyroid organoids formation after lentiviral transduction to silence *Klhl14*, A83-01 (5 mM; Tocris) or equivalent volume of solvent (DMSO) was added to TGM from day 0 of culturing in Basement Membrane Matrigel. The drug was refreshed every three days.

### RNA isolation and qPCR analysis

Total RNA was isolated from thyroid gland organoids using the RNeasy Mini Kit, following the manufacturer’s instructions. RNA was then reverse transcribed using a mix containing 1 µL dNTPs (10 mM), 1 µL random primers (100 ng), 4 µL 5x First-Strand Buffer, 2 µL DTT (0.1 M), 1 µL RNase OUT™ (40 units/µL), and 1 µL M-MLV reverse transcriptase (200 units). Gene expression analysis was performed using specific primers and iQ SYBR Green Supermix, with all reactions carried out in triplicate on a Bio-Rad Real-Time PCR System. The list of primers is provided in Supplementary Table 1, and the Ywhaz gene was used as an internal control.

### Bulk RNA sequencing and analysis

Total RNA was extracted from 5, 7 and 11 days-old organoids as described above. RNA quality was assessed using High Sensitivity RNA ScreenTapes on an Agilent 2100 Bioanalyzer, ensuring highly intact RNA (RIN > 9). RNA samples from organoids derived from three different animals were processed to prepare cDNA libraries using the Smart3SEQ protocol, adhering to the manufacturer’s instructions, with 80 ng of RNA used as input for each sample^41^. The resulting libraries were equimolarly pooled, and 1.8 pM of the pool, supplemented with 15% PhiX, was loaded onto a NextSeq 500 (Illumina) for a 75 bp single-read sequencing run at the Research Sequencing Facility of ERIBA (UMCG). STAR aligner software was used to align the results of the sequencing to mouse genome GRCm39. Z-scores of the genes of interest were handcalculated.

### Protein extraction and western blot analysis

Organoids were harvested after dispase (1 mg/ml) treatment of the gels, lysed in RIPA buffer, sonicated, and centrifuged at 4°C for 5 minutes at maximum speed. The protein concentration of the lysates was determined using the Bradford assay, and the samples were boiled at 99°C for 5 minutes before loading. Equal amounts of protein were resolved on 12% polyacrylamide gels. Proteins were then transferred onto nitrocellulose membranes using the Trans-Blot Turbo System (Bio-Rad). Blocking was carried out with 10% milk in PBS-Tween20 for 1 hour. The membranes were incubated with primary antibodies overnight at 4°C, followed by 1.5-hour incubation at room temperature with horseradish peroxidase-conjugated secondary antibodies, according to the host species of the primaries. The membranes were developed using ECL reagent and visualized with a ChemiDoc imager (Bio-Rad). Western blot analysis was performed using Image Lab software. KLHL14 and GAPDH primary antibodies (Proteintech, 16693-1-AP; Fitzgeral, 10R-G109a) were used respectively with a 1:1000 and 1:10000 dilution in BSA 3% PBS-Tween20; secondary antibodies were used with a 1:3000 dilution in BSA 3% PBS-Tween20.

### Discovery-based proteomic analyses

Discovery-based proteomics analyses were done as described previously (PMID: 37833292). In short, in-gel digestion was performed on the 20 µg total protein lysates with 300 ng trypsin. LC-MS based proteomics analyses were performed injecting an equivalent of 1 µg digested total protein starting material. LC-MS raw data were processed with Spectronaut (version 19.1.240724) (Biognosys) with a Mouse SwissProt database (17141 entries) using the standard settings of the directDIA workflow except that quantification was performed on MS1. For the quantification, the Q-value filtering was set to the classic setting with local normalization and no imputation. For downstream processing Gene set enrichment analysis (GSEA) was performed using the software developed by UC San Diego and Broad Institute^42,43^. Gene ontology terms were obtained through g:Profiler tool^44^.

### Organoids fixation and immunofluorescent staining

To stain organoid sections, thyroid gland organoids were collected and fixed in 4% formaldehyde for 15 minutes, then embedded in HistoGel. The HistoGel-embedded samples were dehydrated, embedded in paraffin, and sectioned into 4 µm thick slices. Sections were dewaxed and subjected to antigen retrieval using Tris-EDTA buffer (pH 9) under heat. After permeabilization and blocking with a solution containing 4% goat serum, 1% BSA, and 0.01% Triton in PBS (1x), the sections were incubated overnight at 4°C with the appropriate primary antibody. Anti-P21 antibody (Abcam, ab188224) 1:100; anti-KLHL14 (Proteintech, 16693-1-AP) 1:200; anti-VIMENTIN (Antibodies.com, A86652) 1:100; anti-LAMIN B1 (Santa Cruz Biotechnology, sc374015) 1:100. The following day, sections were treated with the secondary antibody for 1 hour at room temperature, followed by DAPI staining for 10 minutes to label nuclei. After mounting, images were captured using a Leica DM6 microscope. Image processing was performed with ImageJ (v1.52).

### SA-β-Galactosidase staining

Organoids were harvested on day 7 after lentiviral transduction, fixed, and stained for 4 hours with X-Gal solution (pH 6) following the manufacturer’s instructions (Merck Millipore, KAA002RF). Images of the organoids were captured using a Leica DM6 microscope, and blue-stained cells were identified as senescent cells. The quantification of SA-β-Gal staining was conducted using ImageJ by calculating the ratio of the blue-stained (positive) area to the total organoid area.

### Incucyte Thyroid Organoid Formation Assays

Mouse (human) thyroid organoid cells, after lentiviral transduction, were seeded into 12 (24) multi-well plates and grown in Basement Membrane Matrigel domes as described above. After 7 (21) days in culture, dispase was used to dissolve the gels and 100 ml of each suspension were moved to 96 multi-well plates. The plates were transferred to an IncuCyte® S3 (Essen BioScience) where they imaged using whole-well scans (including phase and green exposure for 300 ms) and kept at 37 °C with 5 % CO_2_ atmosphere. For comparisons between treatments, the total green object integrated intensity was calculated for each well (GCU x µm2/well), and further normalized by the object count per well to estimate average object/organoid sizes.

### Statistical analysis

Statistical analyses were conducted using GraphPad Software version 8. Group differences were evaluated with two-tailed Student’s t-tests or one-way ANOVA with post-hoc Tukey’s HSD. Data are presented as mean ± s.e.m. In all experiments, replicates represent samples from distinct animals, with the number of biological replicates (N) and the statistical test used specified in each figure legend. Statistical significance was defined as *p*-values ≤ 0.05.

### Wound healing assay

Control and shKlhl14 thyroid gland organoids were collected and dissociated into single cells using 0.05% trypsin-EDTA. After cell counting, 50.000 cells/well were re-seeded in a 24-well plate pre-coated with 2% BMM. TGM (± TGFβi) was added, and cells were incubated at 37°C with 5% CO₂. Once cells reached confluency, they were starved by replacing the full medium with DMEM/F12 without growth factors for at least 3 hours. A wound was then created in each well, and plates were then placed in an IncuCyte S3 Live-Cell Analysis System for imaging. Trypan blue was used to assess viability of the non-attached cells from the shKlhl14 DMSO condition.

### 2D culturing of the organoids

7-day-old organoids obtained from transduced cells (shSCRA and shKlhl14) were collected after replacing TGM with dispase (1 mg/ml), reseeded in a 2% BMM-coated 24-well plate then placed in an IncuCyte S3 Live-Cell Analysis System (Essen BioScience) for 96 hours. The wells were imaged using whole-well scans (including phase and green exposure for 300 ms) and kept at 37 °C with 5 % CO_2_ atmosphere. The calculation of the number of organoids with(out) migrating cells was not automatized and averaged on four pictures per well.

## Data availability

The bulk RNA-seq data have been deposited in the Gene Expression Omnibus (GEO) repository; GSE number is available for review purposes and will be made fully available upon acceptance of the manuscript. Proteomic analysis raw data are available as .xlsx file. All other data are stored at the department of Biomedical Sciences, UMCG and are available from the corresponding authors upon request.

## Contributions

R.M. contributed to the conceptualization of the study; acquired, analyzed and interpreted the data; drafted the manuscript. A.A.S-G. acquired and analyzed the data. A.J.J-dB acquired the data. M.E. helped the development of the study. S.K. provided fundings and critical ideas. G.D.V. helped the design of the study and the revision of the manuscript. R.P.C. helped the conceptualization and development of the study, provided fundings, revised the manuscript.

