## Supplemental Material for "Role of Klhl14 in senescence and epithelial-to-mesenchymal transition *via* TGF-β modulation"

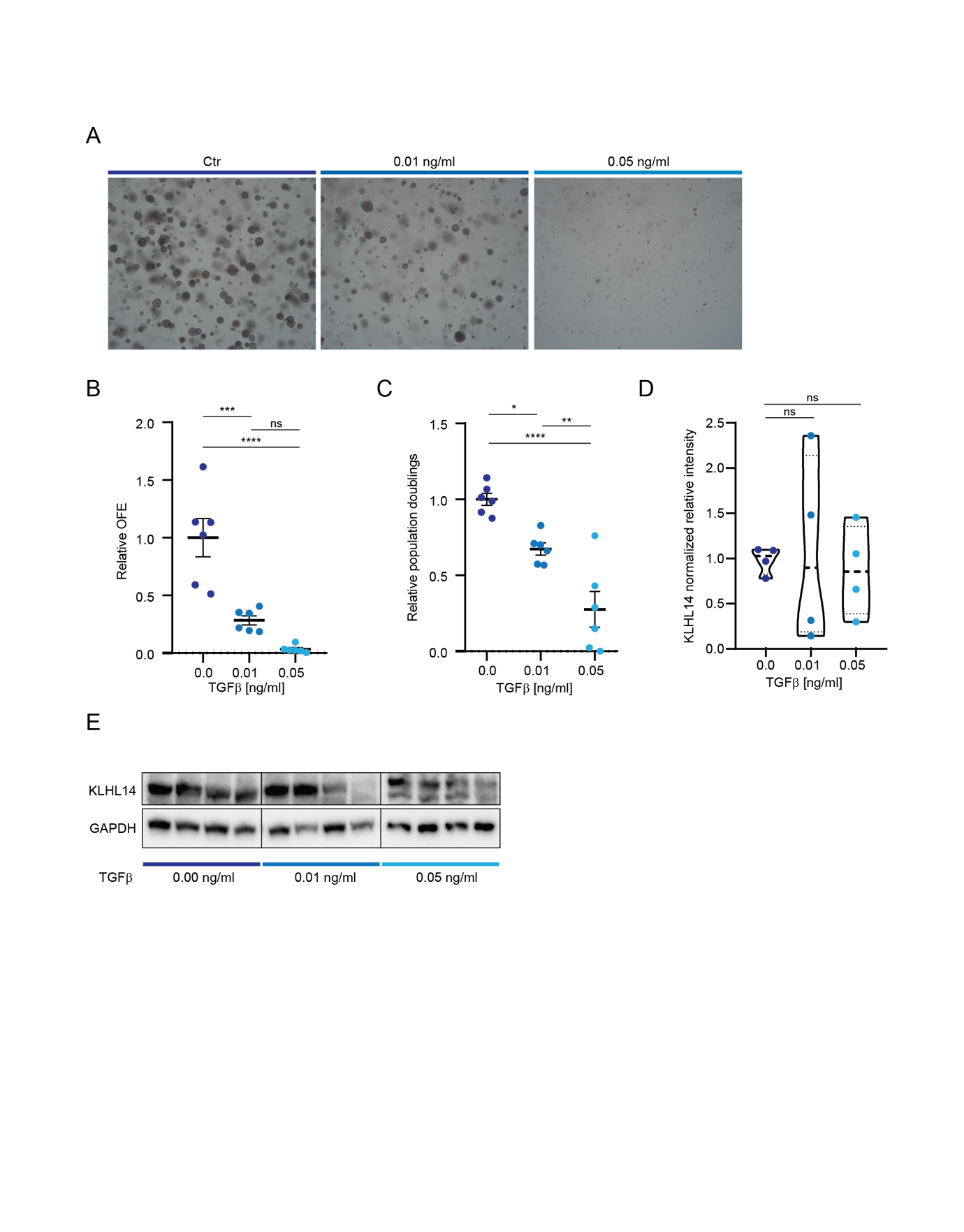

Supplementary Fig. 1: **TGF-β treatments block mTGO growth.** (**a**) Light microscope representative pictures of 7-day-old mTGOs treated with increasing doses of TGF-β, 4x objective. (**b**) Relative OFE calculated on 7-day-old mTGO treated with increasing doses of TGF-β. (**c**) Relative population doublings calculated on 7-day-old mTGO treated with increasing doses of TGF-β. (**d**) Normalized relative intensity of KLHL14 western blot bands reflecting the protein levels in 7-day-old organoids after TGF-β treatment. All the bands in KLHL14 frame have been considered. (**e**) Western blot analysis of 7-day-old organoids grown in presence of TGF-β. Data are shown as mean ± SEM. Group differences were evaluated with one-way ANOVA with post-hoc Tukey’s HSD. N=4-6. **p* < 0.05, ***p* < 0.01, ****p* < 0.001, *****p* <0.0001

| Gene | Primers |
| --- | --- |
| Ywhaz | F: TTACTTGGCCGAGGTTGCT  R: TGCTGTGACTGGTCCACAAT |
| p21 | F: AACATCTCAGGGCCGAAA  R: TGCGCTTGGAGTGATAGAAA |
| p16 | F: GAACTCTTTCGGTCGTACCC  R: CGAATCTGCACCGTAGTTGA |
| Bcl-2 | F: GTGGCCTTCTTTGAG  R: GCCTCCGTTATCCTG |
| Mcp1 | F: GCTCAGCCAGATGCAGTTAA  R: TCTTGAGCTTGGTGACAAAAACT |
| Ccl5 | F: TGCCCACGTCAAGGAGTATTTC  R: AACCCACTTCTTCTCTGGGTTG |
| Klhl14 | F: GGCTCCTCTCCACTCACTTTC  R: TCAGCTCAGCAGCGAAGTC |
| Lmnb1 | F: GGGAAGTTTATTCGCTTGAAGA  R: GGGAAGTTTATTCGCTTGAAGA |
| Cdh1 | F: CGTCCTGCCAATCCTGATGA  R: ACCACTGCCCTCGTAATCGAAC |
| Cdh2 | F: ACAGTGGAGCTCTACAAAGG  R: CTGAGATGGGGTTGATAATG |
| Vim | F: GAACCTCCAGGAGGCCGAGG  R: CATCTTAACATTGAGCAGATC |
| Twist | F: CCAGCTCCAGAGTCTCTAGA  R: GCCAGGTACATCGACTTCCT |
| Zeb1 | F: GCAGAAAATGAGCAAAACCATGA  R: TGGGTTCTGTATGCAAAGGTG |
| Snai1 | F: ATGCACATCCGAAGCCACA  R: GACTCTTGGTGCTTGTGGAG |
| CD24 | F: GTTGCACCGTTTCCCGGTAA  R: CCCCTCTGGTGGTAGCGTTA |
| CD44 | F: TCGATTTGAATGTAACCTGCCG  R: CAGTCCGGGAGATACTGTAGC |
| EpCAM | F: GCGGCTCAGAGAGACTGTG  R: CCAAGCATTTAGACGCCAGTTT |

Supplementary Table 1: **Primers list for qPCR analysis.**
